# Full-length, single-cell RNA-sequencing of human bone marrow subpopulations reveals hidden complexity

**DOI:** 10.1101/2021.07.28.454226

**Authors:** Marcel O. Schmidt, Anne Deslattes Mays, Megan E. Barefoot, Anna T. Riegel, Anton Wellstein

## Abstract

Bone marrow progenitor cell differentiation has frequently been used as a model for studying cellular plasticity and cell-fate decisions. Recent analysis at the level of single-cells has expanded knowledge of the transcriptional landscape of human hematopoietic cell lineages. Using single-molecule real-time (SMRT) full-length RNA sequencing, we have previously shown that human bone marrow lineage-negative (Lin-neg) cell populations contain a surprisingly diverse set of mRNA isoforms. Here, we report from single cell, full-length RNA sequencing that this diversity is also reflected at the single-cell level. From fresh human bone marrow unselected and lineage-negative progenitor cells were isolated by droplet-based single-cell selection (10xGenomics). The single cell-derived mRNAs were analyzed by full-length SMRT and short-read sequencing. In both samples we detected an average of 8000 different genes using short-read sequencing. Differential expression analysis arranged the single-cells of the total bone marrow into only four clusters whereas the Lin-neg population was much more diverse with nine clusters. mRNA isoform analysis of the single-cell populations using full-length sequencing revealed that Lin-neg cells contain on average 24% more novel splice variants than the total bone marrow cells. Interestingly, among the most frequent genes expressing novel isoforms were members of the spliceosome, e.g. HNRNPs, DEAD box helicases and SRSFs. Mapping the isoforms from all genes to the cell type clusters revealed that total bone marrow cells express novel isoforms only in a small subset of clusters. On the other hand, lineage-negative progenitor cells expressing novel isoforms were present in nearly all subpopulations. In conclusion, on a single-cell level lineage-negative cells express a higher diversity of genes and more alternatively spliced novel isoforms suggesting that cells in this subpopulation are poised for different fates.

**Graphical abstract:** 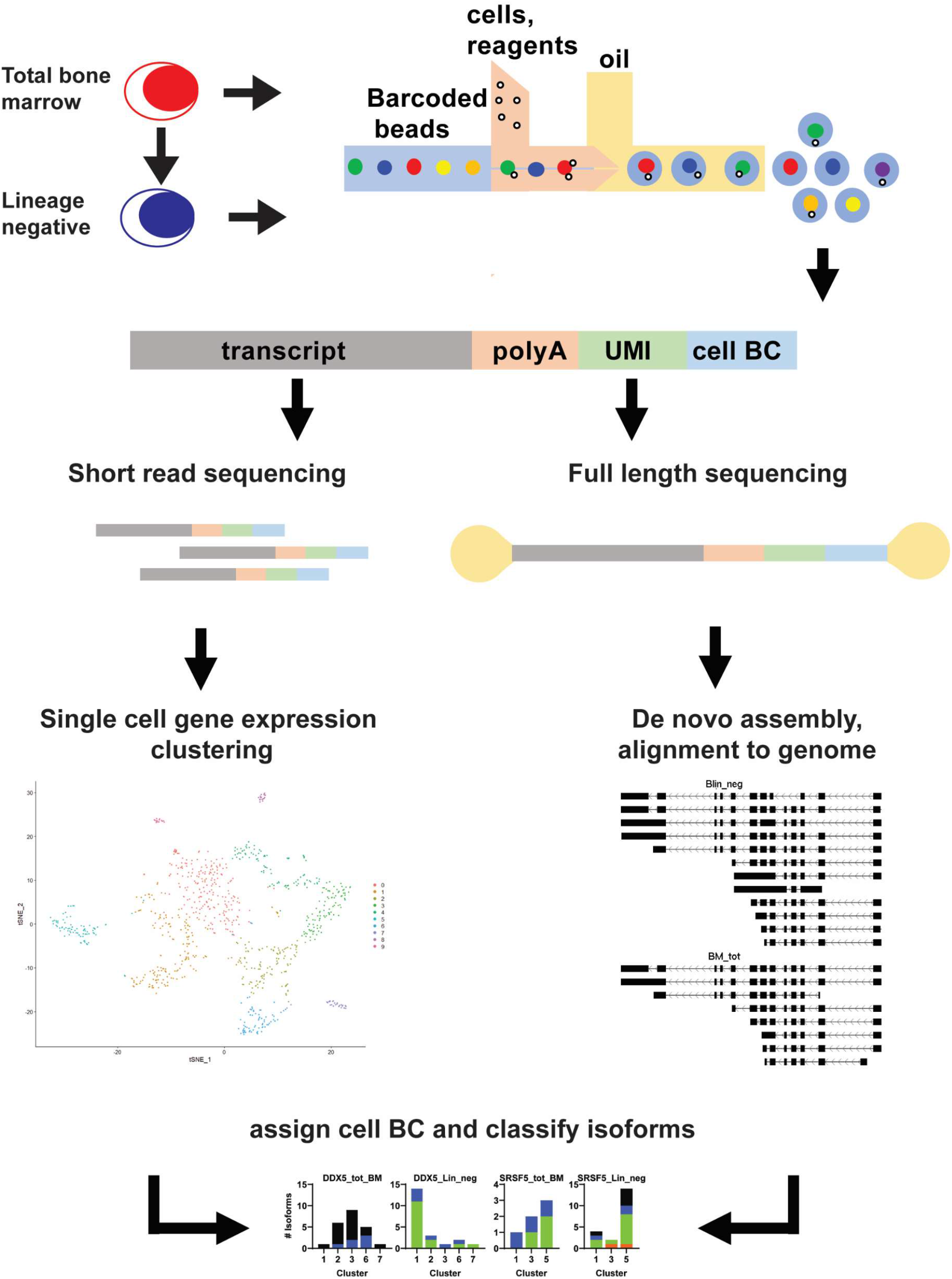

## Introduction

In 2001, the International Human Genome Sequencing Consortium published that a typical human gene contains nine exons (Consortium 2004). During posttranscriptional editing of the nascent transcripts the introns are removed and exons spliced together to generate the coding or messenger RNA (mRNA). Alternative splicing of exons increases the diversity of mRNA isoforms, facilitating the generation of a multitude of protein variants with divergent functions from a single gene locus. Well-known examples for alternatively spliced gene products with distinct functions include isoforms of the *FAS* and *BCL-X* genes with either pro- or anti-apoptotic activities (Cheng et al. 1994; Boise et al. 1993). Also, the pro-angiogenic *VEGF* gene product can be alternatively spliced to produce an isoform with anti-angiogenic function (Nowak et al. 2008). In addition, multiple transcript isoforms are generated from the *TP53* tumor suppressor gene that are differently expressed between cancer and normal tissues (Vieler and Sanyal 2018).

Over the last decade technological advances have enabled gene expression analysis at the single-cell level. This has led to expanded knowledge of cellular diversity in terms of gene-expression. However, only few studies have aimed to characterize this diversity of splice variants at the single cell level (reviewed in Hardwick et al. 2019; Arzalluz-Luque and Conesa 2018). Several studies have evaluated isoform expression from single cells in mouse brain (Karlsson et al. 2017; Gupta et al. 2018) and in mouse B cells (Byrne et al. 2017). Nonetheless, similar studies have yet to be conducted with single-cells from human bone marrow, despite it being a continuously differentiating tissue that contains distinct subpopulations, and has a high turnover rate. The single-cell gene expression levels of human bone marrow-derived hematopoietic stem/progenitor cells have been described by several laboratories (Pellin et al. 2019; Velten et al. 2017). We have found that lineage-negative progenitor cells isolated from bulk bone marrow express a high diversity of transcript isoforms including many novel splice variants not previously known (Deslattes Mays et al. 2019). Here, we investigate the extent to which individual cells within the total bone marrow (tot-BM) and the lineage negative (Lin-neg) populations exhibit this isoform diversity and whether distinct subpopulations in tot-BM and Lin-neg cells intersect based on transcript isoform usage. Using a droplet-based approach, we separately isolate tot-BM cells and Lin-neg cells, and sequence unfragmented libraries of full-length mRNA using Single Molecule Real-Time (SMRT) RNAseq technology on the PacBio platform.

## Results

We extracted healthy donor human bone marrow tissues from discarded harvesting filters. From this total bone marrow cell preparation (tot-BM) we enriched for lineage-negative progenitor cells (Lin-neg) by magnetic selection as described earlier (Deslattes Mays et al. 2019). We then analyzed total and Lin-neg cell populations by droplet-based single cell RNA sequencing (10xGenomics). In order to increase the number of mRNA molecules detectable per cell we reduced the cell input to approximately 500 cells for each experiment. After single-cell selection, the barcoded cDNA libraries were equally divided and each pool was analyzed in parallel by short-read sequencing (Illumina) and by single-molecule real-time (SMRT) full-length RNA sequencing (Pacific Biosciences). The short-read sequencing results were used to cluster the single cells based on gene expression and identify the different cell types present in each sample. Full-length mRNA sequencing results were used to identify the transcript isoforms present within individual cells in different clusters.

### Short-read sequencing reveals greater diversity in lineage-negative subpopulation compared to total bone marrow populations

The 10X-Genomics’ microfluidic device isolates each cell in a water/oil emulsion resulting in cell-based and barcode tagged libraries with each cDNA molecule containing a unique molecular identifier (UMI). We split the cDNA libraries into two equal pools, one of which was analyzed by short-read RNAseq of fragmented cDNAs (Illumina) and the other by SMRT full-length RNAseq (PacBio).

The short-read data was evaluated with the Seurat package and we detected 415 total BM (tot-BM) and 492 lineage-negative (Lin-neg) cells. While we found comparable gene numbers in each sample (7212 genes in tot-BM, 8758 in Lin-neg) the number of distinct mRNA molecules detected using the UMI varied by over two-fold, i.e. 57,322 for tot-BM and 133,744 for Lin-neg (Table 1). We clustered the cells based on their gene expression levels and identified five clusters in tot-BM and nine in Lin-neg cell populations. We identified the most differentially expressed genes in each cluster and annotated each cluster by cell type (Fig. 1, Table 2). Supplemental. Tables S1 and S2 show the top marker genes expressed in each cluster. Interestingly, in addition to the five clusters identified from tot-BM cells we found four more clusters in the Lin-neg cells. Additional clusters were CD34+ progenitor (cluster 3), early B-cells (clusters 6,8), and immature granulocytes and neutrophils (cluster 2).

**Table 1:** Total number of genes and mRNA molecules (UMI) from short-read data in single cells

| sample | genes | UMI |
| --- | --- | --- |
| BM_tot | 7,212 | 57,322 |
| Blin_neg | 8,758 | 133,744 |

**Table 2:**
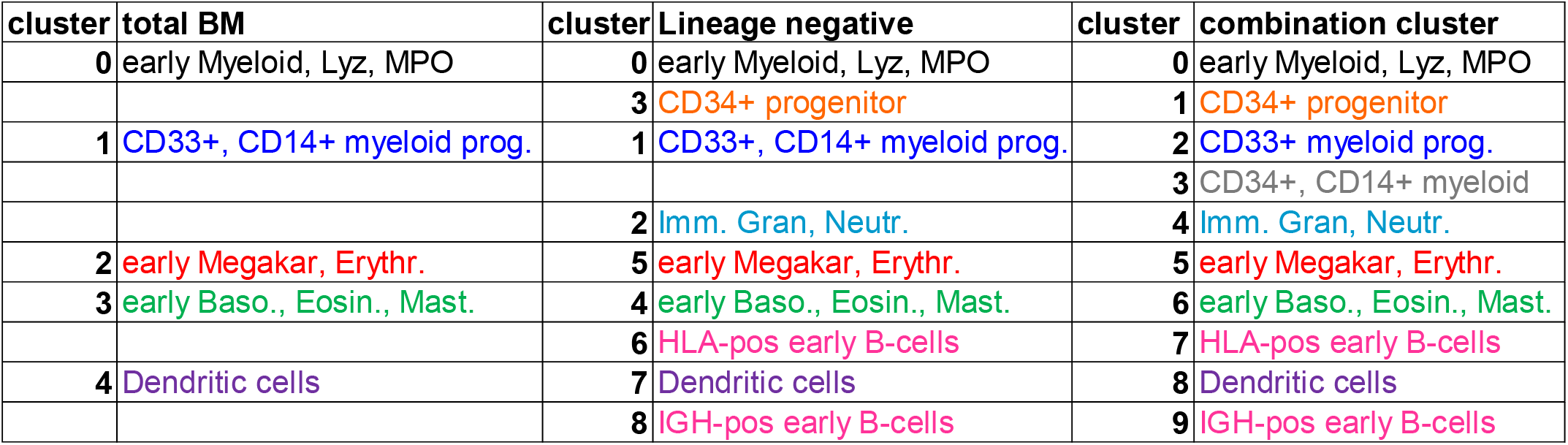
Cluster gene type annotation by expression aligned by row (BioGPS, The Human Protein Atlas, Pellin et al. 2019, Velten et al. 2017)

| cluster | total BM | cluster | Lineage negative | cluster | combination cluster |
| --- | --- | --- | --- | --- | --- |
| 0 | early Myeloid, Lyz, MPO | 0 | early Myeloid, Lyz, MPO | 0 | early Myeloid, Lyz, MPO |
|  |  | 3 | CD34+ progenitor | 1 | CD34+ progenitor |
| 1 | CD33+, CD14+ myeloid prog. | 1 | CD33+, CD14+ myeloid prog. | 2 | CD33+ myeloid prog. |
|  |  |  |  | 3 | CD34+, CD14+ myeloid |
|  |  | 2 | Imm. Gran, Neutr. | 4 | Imm. Gran, Neutr. |
| 2 | early Megakar, Erythr. | 5 | early Megakar, Erythr. | 5 | early Megakar, Erythr. |
| 3 | early Baso., Eosin., Mast. | 4 | early Baso., Eosin., Mast. | 6 | early Baso., Eosin., Mast. |
|  |  | 6 | HLA-pos early B-cells | 7 | HLA-pos early B-cells |
| 4 | Dendritic cells | 7 | Dendritic cells | 8 | Dendritic cells |
|  |  | 8 | IGH-pos early B-cells | 9 | IGH-pos early B-cells |

**Figure 1.**
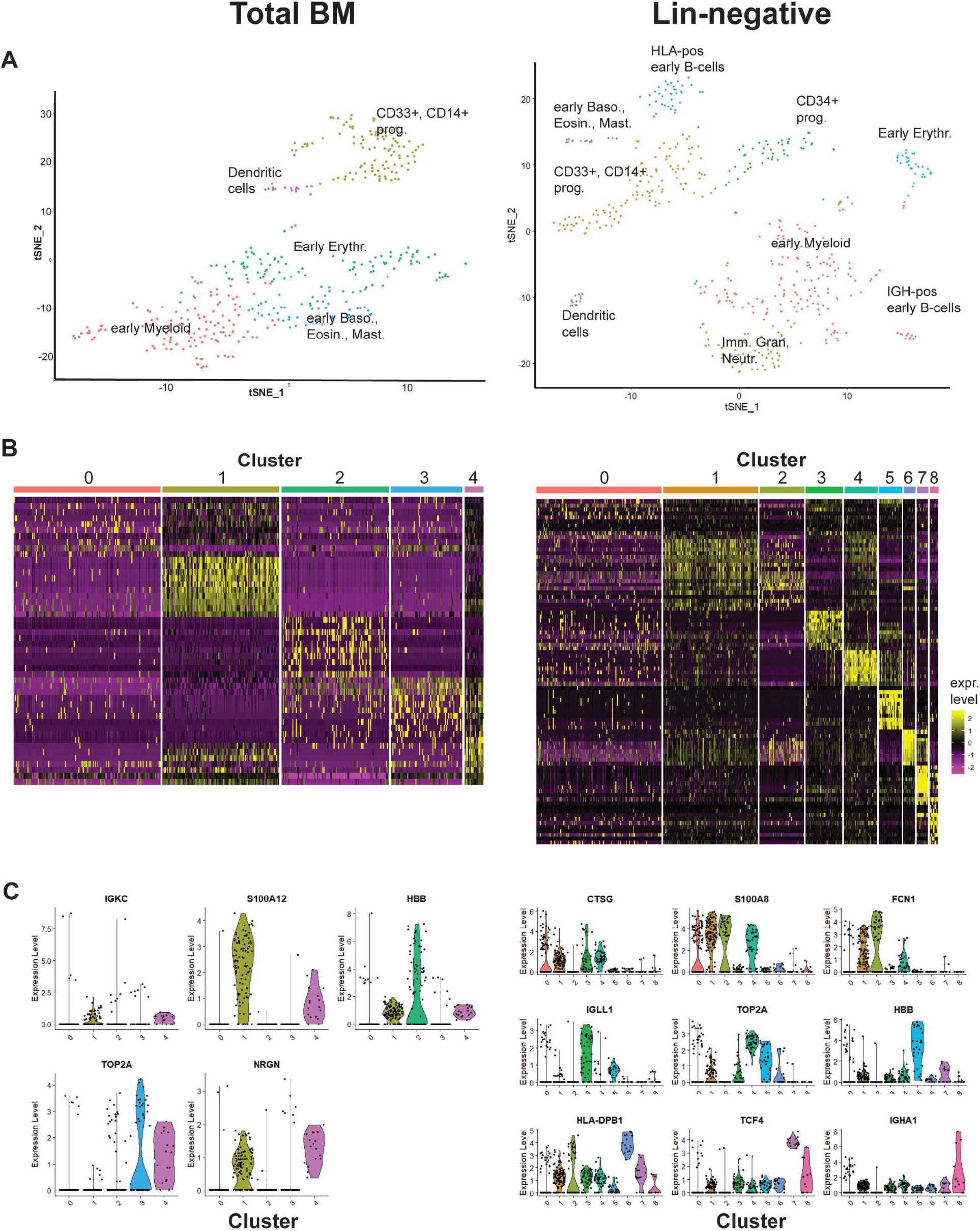
Lineage-negative cells have greater diversity than total bone marrow cells. (**A**) Single-cell short-read expression analysis with the Seurat R package reveal 5 clusters for tot-BM cells (left panel) and 9 for Lin-neg cells (right panel). Clusters are annotated according to marker gene expression (Wu et al. 2016; Uhlén et al. 2015; Pellin et al. 2019; Velten et al. 2017). (**B**) Heatmaps of the top 10 regulated genes for each cluster. (**C**) Violin expression plots of the top cell type marker for each cluster. The full names of genes indicated as acronyms are listed in Suppl. Tables S1 and S2

To identify potential overlap of clusters, we combined both samples in-silico and re-clustered the samples. The re-clustered cells separated into 10 clusters and we annotated the cell types in each cluster (Fig. 2A, Table 2). Supplemental Table S3 and Supplemental Fig. S1 show the top marker genes defining these cell type clusters. The combination analysis revealed one additional cluster 3 with more mature CD34, CD14 positive myeloid cells, which was populated by a majority of tot-BM cells. Also, cluster 5 with early megakaryocytes contained more tot-BM cells (Fig. 2). In comparison, Lin-neg cells were the majority contributors to two distinct populations of early B-cells identified from the combination clustering analysis, expressing either immunoglobulin genes (cluster 7) or MHC II (HLA) genes (cluster 9). The majority of clusters contain more Lin-neg than tot-BM cells reflecting the diversity of Lin-neg cells (Fig. 2B, Table 3).

**Table 3:** Number of cells in each cluster after filtering data with Seurat package. The distribution of cells into clusters is significantly different between tot-BM and Lin-neg cell populations (P<0.0001; chi-square test)

| cluster | tot-BM | lin.neg | all |  |
| --- | --- | --- | --- | --- |
| 0 | 95 | 118 | **213** | chi-square test: $p < 0.0001$ |
| 1 | 79 | 93 | **172** |  |
| 2 | 40 | 80 | **120** |  |
| 3 | 64 | 33 | **97** |  |
| 4 | 39 | 54 | **93** |  |
| 5 | 45 | 35 | **80** |  |
| 6 | 32 | 40 | **72** |  |
| 7 | 13 | 14 | **27** |  |
| 8 | 4 | 14 | **18** |  |
| 9 | 4 | 11 | **15** |  |
| **Total** | **415** | **492** | **907** |  |
| | chi-square test:<br>$p < 0.0001$ | | | |

**Figure 2.**
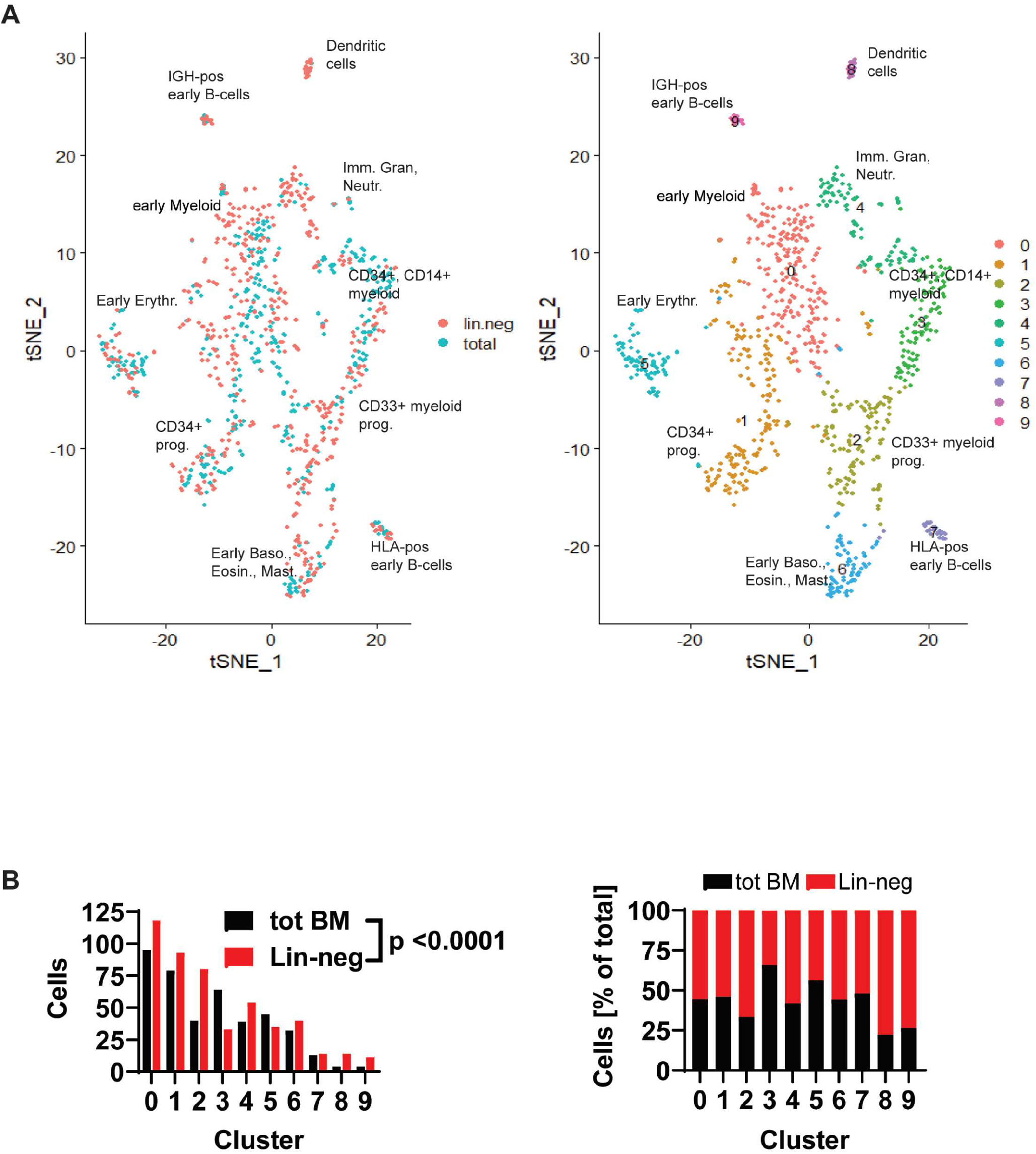
Most clusters of the combined single-cell expression analysis contain more Lin-neg than tot-BM cells. (**A**) tSNE plot of combined cluster analysis colored by sample (left panel) or by cluster (right panel). Clusters are annotated according to marker gene expression (Wu et al. 2016; Uhlén et al. 2015; Pellin et al. 2019; Velten et al. 2017). (**B**) Total number (left) and fraction of cells per cluster (right). Chi-square tests for total BM and Lin-neg cells number distribution in clusters show highly significant distributions (p < 0.0001)

### Full-length RNA sequencing reveals higher rate of novel isoforms in Lin-neg cell populations

Using the parallel pool of barcoded and UMI labeled single-cell cDNA libraries, we performed full-length SMRT sequencing that avoids fragmentation of the cDNA. Using the scalable de novo isoform discovery workflow, Isoseq3, we detected approximately 40 million circular consensus reads. We required three read passes to create a high-fidelity consensus sequence. Furthermore, we filtered the consensus reads to contain both 3’ and 5’ adapters, cell barcodes, UMI sequences and poly A tails to ensure that the sequences represent full-length molecules. The de-novo assembly generated 260,000 and 340,000 transcripts for the tot-BM and Lin-neg samples, respectively (Table 4). The full-length reads were then mapped to the reference transcriptome using the Isoseq3 and cDNA cupcake packages. Transcript isoforms were annotated using the SQANTI3 package filtering out intra-priming instances. We found transcripts from 7,670 and 9,720 genes and 14,781 and 23,101 unique transcript isoforms in tot-BM and Lin-neg, respectively (Table 4). The transcript isoforms were classified into four categories with the SQANTI3 package (Tardaguila et al. 2018): The “Full Splice Match” (FSM) indicates an exact match with the number of exons and splice junctions of the reference transcriptome; isoforms in the “Incomplete Splice Match” (ISM) category lack 5’ and/or 3’ exons; “Novel In Catalog” (NIC) isoforms use new combinations of known junctions and “Novel Not In Catalog” (NNIC) are novel isoforms with at least one new splice junction (Fig. 3A).

**Table 4.** Full-length sequencing workflow using CCS, lima, isoseq3, minimap2 and SQANTI

|  | **tot-BM** | **Lin-neg** |
| --- | --- | --- |
| raw reads | 39,270,949 | 44,507,070 |
| **CCS --minPasses 3** | 393,537 | 444,826 |
| **lima** require 5p to 3p adapters | 274,879 | 365,226 |
| clip_out_UMI_cellBC.py | 270,545 | 358,406 |
| **isoseq3** |  |  |
| num_reads_flnc | 258,783 | 347,801 |
| num_reads_flnc_polya | 256,155 | 342,805 |
| **minimap2** |  |  |
| mapped sequences | 256,155 | 342,805 |
| **sqanti "flnc"** |  |  |
| unique genes | 18,965 | 24,110 |
| unique isoforms | 66,820 | 104,728 |
| **sqanti_filtered** |  |  |
| unique genes | 7,670 | 9,720 |
| unique isoforms | 14,781 | 23,101 |

**Figure 3.**
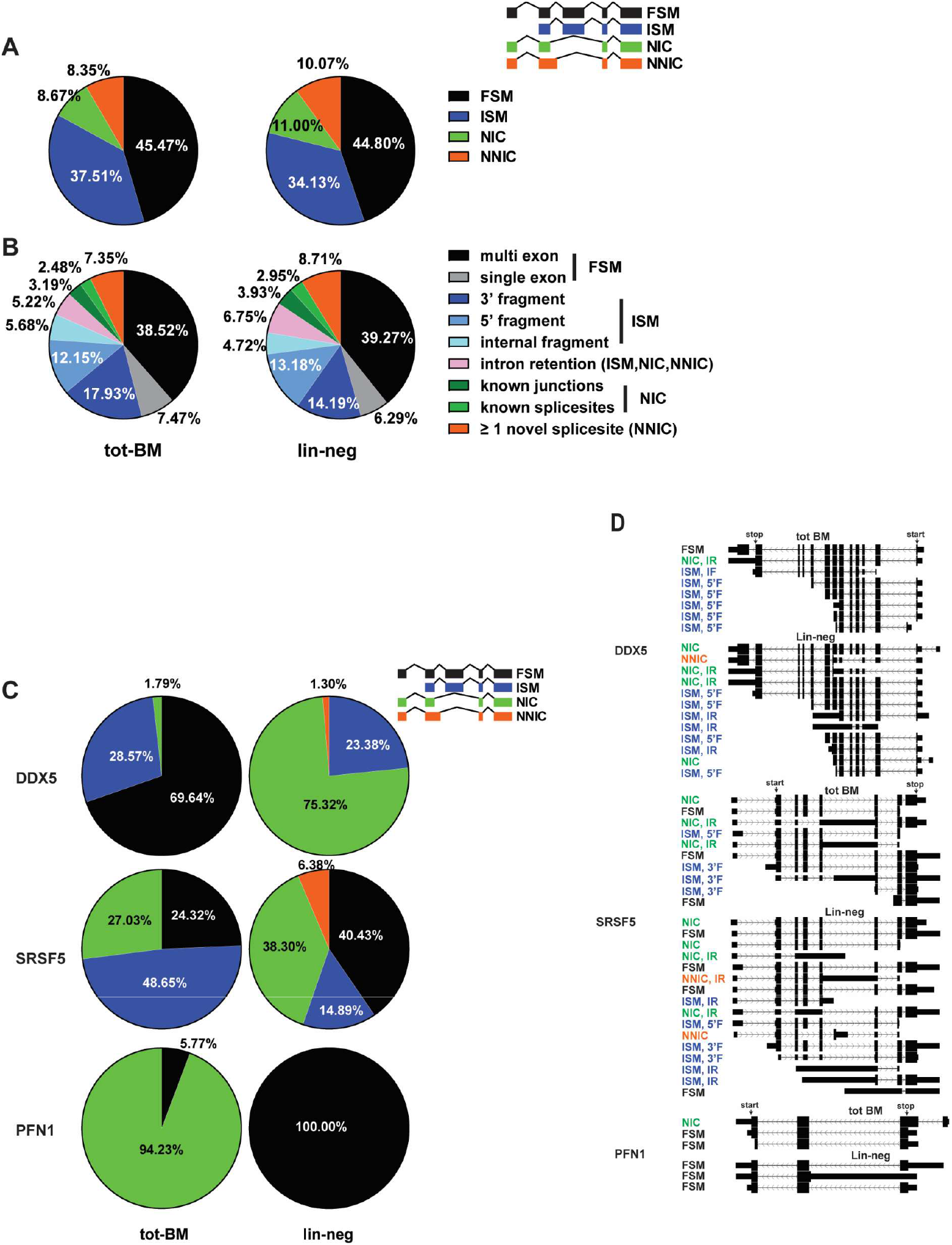
Lin-neg cells express more novel isoforms than tot-BM cells. (**A**) Major isoform categories for each sample. Consensus isoforms are either “Full Splice Match” (FSM) or “Incomplete Splice Match” (ISM). Novel isoforms are either “Novel In Catalog” (NIC) or “Novel Not In Catalog” (NNIC). (**B**) Subcategories of alternatively spliced isoforms. Multi and single exon are subcategories of FSM. 3’ fragment, 5’ fragment, and internal fragment are subcategories of ISM. The NIC category is subdivided into known junctions and known splice sites. NNIC contains novel splice sites. ISM, NIC and NNC categories may also contain an intron retention. (**C**) Isoform categories and relative content for DDX5, SRSF5 and PFN1. (**D**) UCSC genome browser image of unique isoforms detected in each sample. Isoforms are labeled with categories and subcategories. Multi exon (ME), single exon (SE), 3’ fragment (3’F), 5’ fragment (5’F), internal fragment (IF), known junctions (KJ), known splice sites (KS), intron retention (IR). Consensus translation starts and stops are indicated. Chi-square tests for total BM and Lin-neg isoform distribution show highly significant distributions (p < 0.0001) for A, B and C (DDX5, PFN1), p=0.0054 for SRSF5)

Remarkedly, in the Lin-neg cells, we observed a significant 27% increase (from 8.7% (tot-BM) to 11% (Lin-neg)) in splice variants with novel combinations of known exons (NIC) as well as a 21% increase (from 8.35% to 10.07%) isoforms with at least one novel splice site (NNIC) (Fig. 3A). Further subtyping of each isoform is shown in Fig. 3B and Table 5. FSM is further subcategorized into single-(SE) and multi-exonic (ME) genes. ISM is subclassified as internal (IF), as 5’ or 3’ fragments (5’F, 3’F). NIC contains combinations of known splice sites (KS) or junctions (KJ). In addition, ISM, NIC and NNC categories are subcategorized as having an intron retention (IR) or not. Overall, Lin-neg cells are 29% more enriched for novel isoforms (both NIC and NNIC) with both known and novel junctions and splice sites. This includes enrichment of splice variants containing intron retentions (IR, from 5.22% to 6.75%), which can lead to nonsense-mediated decay (NMD) (Jacob and Smith 2017).

**Table 5.** Alternative splicing events as categorized by SQANTI are significantly different between tot-BM and Lin-neg cell populations (P<0.0001; chi-square test)

| structural_category | subcategory | tot-BM [#] | tot-BM [%] | Lin_neg [#] | Lin_neg [%] | chi-square test: p<0.0001 |
| --- | --- | --- | --- | --- | --- | --- |
| full-splice_match | multi-exon | 5160 | 38.52% | 8063 | 39.27% |  |
| full-splice_match | single-exon | 1001 | 7.47% | 1291 | 6.29% |  |
| incomplete-splice_match | 3prime fragment | 2402 | 17.93% | 2913 | 14.19% |  |
| incomplete-splice_match | 5prime fragment | 1627 | 12.15% | 2706 | 13.18% |  |
| incomplete-splice_match | internal fragment | 761 | 5.68% | 970 | 4.72% |  |
| ISM, NIC, NNIC | intron retention | 699 | 5.22% | 1385 | 6.75% |  |
| novel_in_catalog | combination_of_known_junctions | 427 | 3.19% | 807 | 3.93% |  |
| novel_in_catalog | combination_of_known_splicesites | 332 | 2.48% | 606 | 2.95% |  |
| novel_not_in_catalog | at_least_one_novel_splicesite | 985 | 7.35% | 1789 | 8.71% |  |

### Genes involved in mRNA processing show a high frequency of novel isoforms

Observing an overall increase of novel isoforms in the Lin-neg population led us to identify individual genes expressing novel variants. For that, we filtered the genes for expression levels (≥20 full-length molecules) and sorted them by the highest ratio for novel isoforms for tot-BM and Lin-neg cells (Table 6). Surprisingly, we found among both populations novel isoforms of genes involved in RNA splicing, such as members of the Heterogeneous Nuclear Ribonucleoproteins (HNRNPs), DEAD-Box Helicase 5 (DDX5, Fig. 3 C, D), Serine And Arginine Rich Splicing Factor 5 (SRSF5, Fig. 3 C, D), Splicing Factor 1 (SF1), Ribonuclease T2 (RNASET2), and a Pre-mRNA Processing Factor (PRPF40A). While both samples revealed novel HNRNP isoforms, novel splice variants of DDX5, PRPF40A, RNASET2 and SRSF5 were enriched in the Lin-neg sample. On the other hand, novel splice isoforms of SF1 predominated in tot-BM.

**Table 6.**
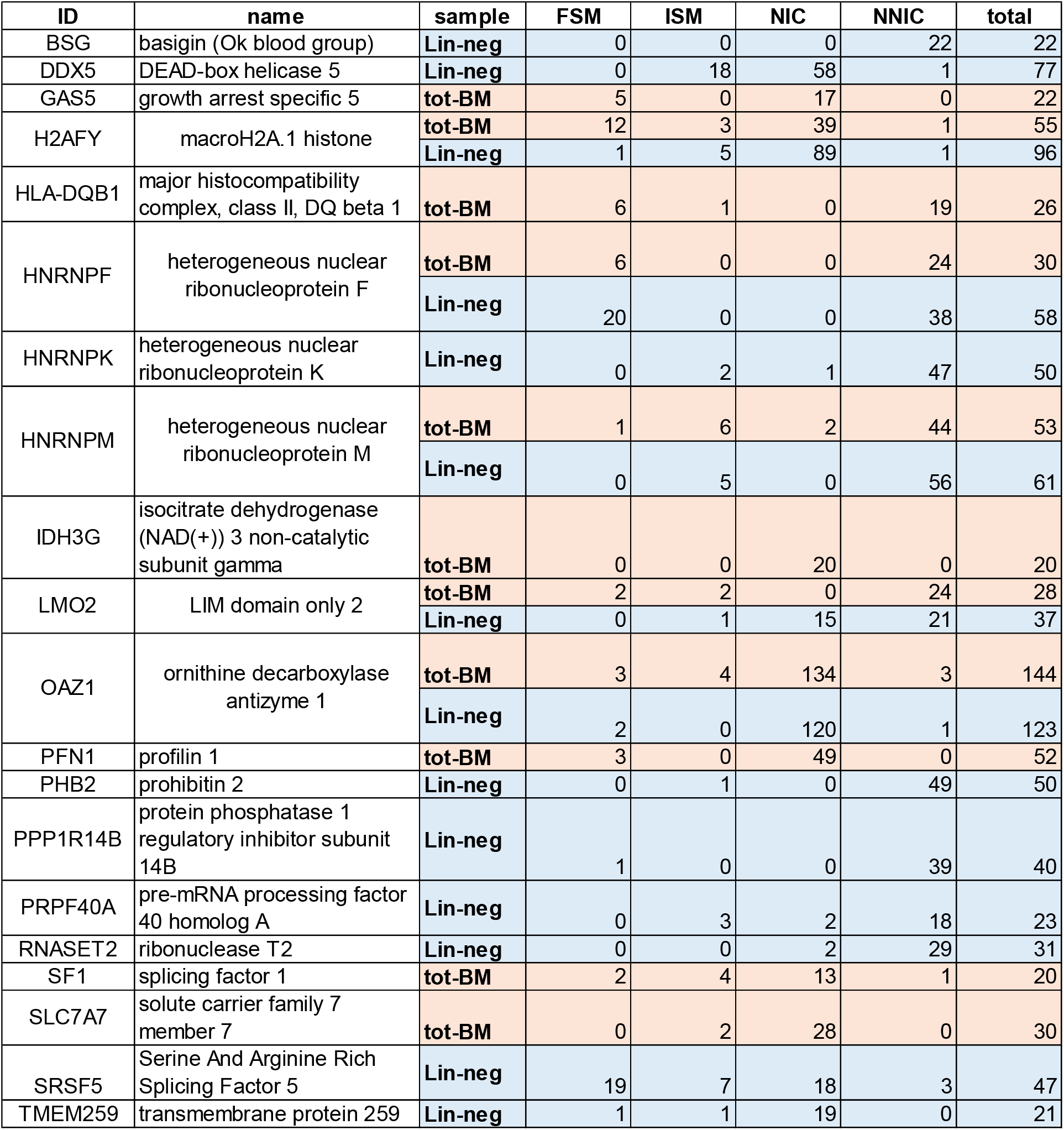
Genes with the most novel isoforms in tot-BM and Lin-neg samples (FSM: Full Splice Match, ISM: Incomplete Splice Match, NIC: Novel in Catalog, NNIC: Novel not in Catalog)

Among other genes expressing novel splice variants, we found the Ornithine Decarboxylase Antizyme 1 (OAZ1, involved in polyamine metabolism), LIM Domain Only 2 (LMO2) maintains pre-erythrocytes in immature state) and a histone-encoding gene (H2AFY). Only few genes expressed more novel isoforms in the tot-BM than in the Lin-neg sample. Among them were Profilin 1 (PFN1) that is involved in cytoskeleton maintenance (Fig. 3 C, D), a member of the Major histocompatibility class II complex (HLA-DQB1) and an isocitrate dehydrogenase (IDH3G) involved in the citrate cycle.

From our cluster analysis, we found several differentially expressed alternative splicing-associated genes in Lin-neg versus tot-BM cells (Fig. 4). Splicing factors of the DDX helicase(Bourgeois et al. 2016) (e.g. DDX5) and the SRSF family (Twyffels et al. 2011) (e.g. SRSF5 and SRSF7) are higher expressed by Lin-neg cells in clusters 1, 2, and 5 to 9 while downregulated in cluster 4. Similarly, the PI3K related kinase (SMG1) (Chen et al. 2017), is also upregulated in clusters 1,2, and 5 to 8 while downregulated in cluster 4. Another gene involved in mRNA processing such as HNRNPA1 (Chen et al. 2010) had a similar expression pattern (Fig. 4). Of particular interest, SMG1 is involved in nonsense-mediated mRNA decay (NMD, (Powers et al. 2020)) and we have observed an overall increase of intron retention in Lin-neg cells (Fig. 3B). This observation led us to investigate the expression of “poison exons” of SRSF genes, defined as ultra-conserved regions containing noncoding exons, that, cause NMD when spliced-in (Lareau et al. 2007b; Leclair et al. 2020; García-Moreno and Romão 2020; Jacob and Smith 2017). Indeed, we found such sequences for SRSF2, 5, 6 and 7 when we entered the coordinates published in (Leclair et al. 2020). Expression of the poison exons were increased in the Lin-neg population. Lin-neg cells express more poison exons of SRSF1,3,5,6, and 7 than tot-BM cells. Only poison exons of SRSF2 and 4 are expressed equally between the populations (Supplemental Fig. S2).

**Figure 4.**
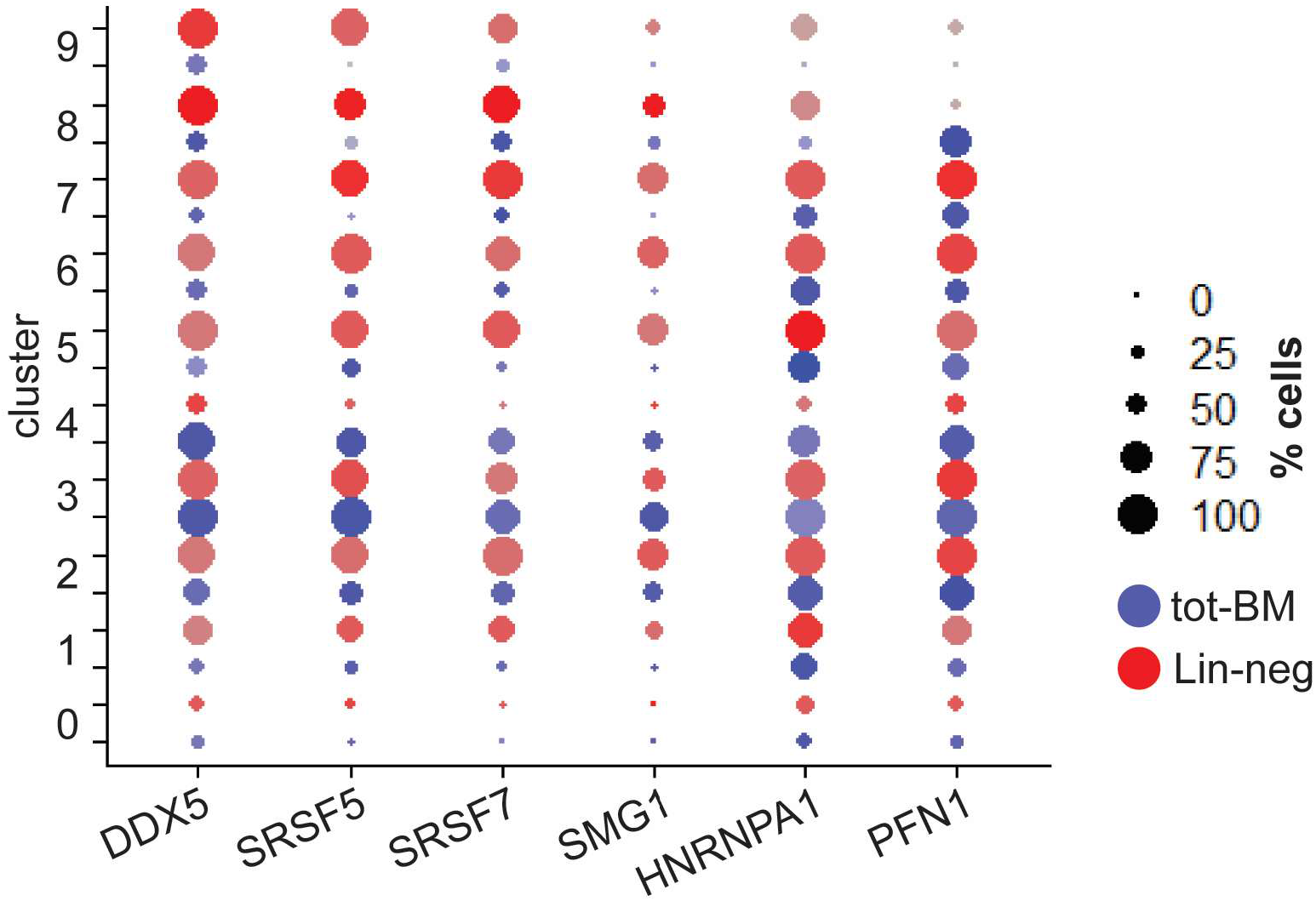
Spliceosome-associated genes are alternatively spliced and have higher expression in Lin-neg cells in most clusters Dot plots of expression levels of alternatively spliced genes detected by short-read sequencing arranged in clusters. Size of dots corresponds to percentage of cells expressing the gene, Color tint reflects average expression level.

### Lin-neg cells with novel isoforms map to most cell type clusters

After identification of alternative splicing of isoforms by full-length RNA sequencing we then used the single-cell barcodes to assign them to cell type clusters classified by short-read RNA sequencing. This classification reduced the total transcript numbers to 34,000 and 54,000 full-length transcripts for tot-BM and Lin-neg, respectively. (Table 7). We then filtered for cells with at least five isoforms. Those isoforms were expressed in 144 tot-BM and 292 Lin-neg cells. We mapped these cells to the combined cell type clusters and found that tot-BM cells with full-length isoforms spread across a subset of clusters (Fig. 5A, left panel). Clusters with CD33+ myeloid progenitor cells (cluster 2), CD34+/CD14 myeloid cells (cluster 3), and immature granulocytes/neutrophils (cluster 4) were the clusters containing the most selected cells. Only 4 full-length transcripts of tot-BM cells were detected in cluster 0 containing early myeloid cells. In contrast, Lin-neg cells were more evenly distributed among the clusters. Only clusters 0 and 4 were somewhat underrepresented (Fig. 5A, right panel). We then sorted the cells by isoform classification (FSM, ISM, NIC, and NNIC) and mapped the top 25 cells in each category to the cell type clusters (Fig. 5B). We found that Lin-neg cells containing all isoform categories were widely distributed in nearly all clusters. Clusters with dendritic (cluster 8), IGH-pos early B-cells (cluster 9) and early erythrocytes (cluster 5) did not contain Lin-neg cells with a majority of full splice matches. In contrast, novel isoforms (ISM, NIC, and NNIC) were found in all clusters except in cluster 0 with early myeloid cells and cluster 4 with immature granulocytes/neutrophils (Fig. 5B, bottom panel). This distribution pattern was in stark contrast to the tot-BM cells, which we found to be enriched only in a smaller subset of clusters. Tot-BM cells with a majority of reference matching isoforms (FSM and ISM) were observed in clusters 1, 2, 3, 5, 6, and 8, while cells with the highest content of novel isoforms were mostly in clusters 2, 3, and 4 (Fig. 5B, top panel). We then compared the isoform expressions of the tot-BM and Lin-neg samples in all cells for each cluster directly (Fig. 5C). In a subset of clusters, Lin-neg cells with novel isoforms were enriched relative to tot-BM cells. We found that cells in clusters 5, 7, 8, and 9 contained novel isoforms from the Lin-neg population (NIC, or NNIC) whereas tot-BM cells with novel isoforms were not significantly increased. Interestingly, in cluster 0 (early Myeloid cells) with a total of 213 cells, only 89 and 4 full-length isoforms of Lin-neg and tot-BM populations, respectively, were detected. (Table 3, 7). Overall, we conclude that Lin-neg cells more frequently express a greater diversity of novel isoforms in a subset of gene expression clusters.

**Table 7.**
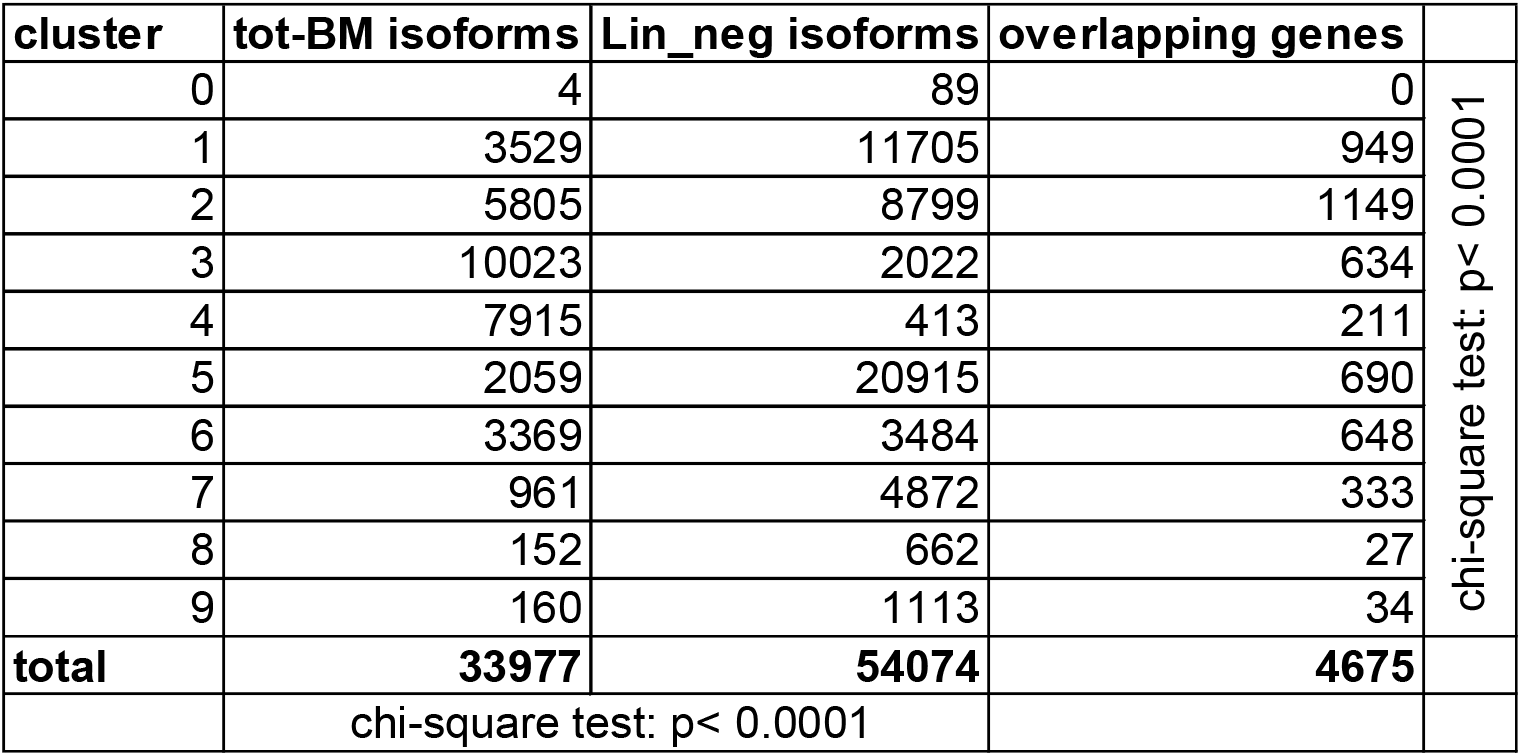
Isoforms per cluster. The distribution of isoforms in the different clusters is significantly different between tot-BM and Lin-neg cell populations (P<0.0001; chi-square test).

**Figure 5.**
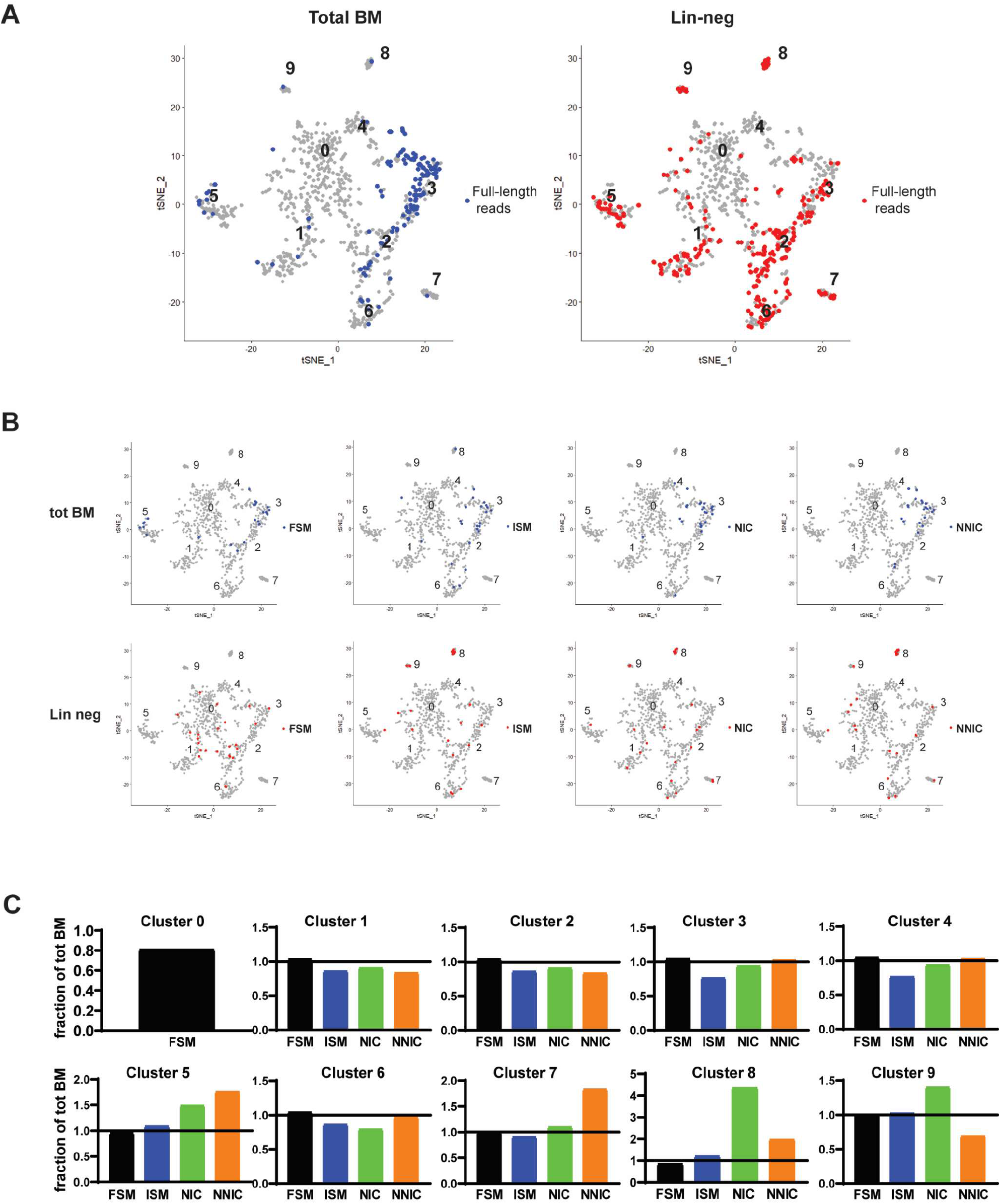
Lin-neg cells expressing novel isoforms are found in most clusters. (**A**) tSNE plot of tot-BM cells (left) and Lin-neg cells (right). Cells containing full-length reads filtered for at least 5 reads per gene are highlighted in blue (tot-BM) and in red (Lin-neg). (**B**) tSNE plots depict the top 25 cells of each isoform category of the tot-BM (highlighted in blue, top panels) and the Lin-neg population (highlighted in red, bottom panels). (**C**) Bar graph showing expression of isoform categories in Lin-neg cells by cell type cluster. Full Splice Match = FSM, Incomplete Splice Match = ISM, Novel In Catalog = NIC, Novel Not In Catalog = NNIC.

### mRNA processing genes have the highest frequency of novel isoforms and are expressed across multiple cell type clusters

Next, we searched for cluster-specific splice variants of single genes. DDX5, the helicase involved in RNA splicing, expressed in all clusters (see Fig. 4) is highly alternatively spliced in Lin-neg cells (see Fig. 3C, D) and we found full-length transcripts in clusters 1,2,3,6, and 7. Interestingly tot-BM cells express canonical isoforms whilst Lin-neg cell isoforms were mostly novel (Fig. 6A). Isoforms of SRSF5, which is also expressed in most clusters and differentially spliced in the Lin-neg population (see Fig. 4 and 3C, D), were detected in clusters 1, 3, and 5 (Fig. 6A). Notably, tot-BM expressed only very few SRSF5 molecules (1 to 3) half of which were novel transcript isoforms. In contrast, Lin-neg cells contained more transcripts (2 to 14), of which the majority were novel (NIC or NNIC) (Fig. 6A). Cells with transcripts isoforms of the Heterogeneous Nuclear Ribonucleoproteins (HNRNPM and HNRNPF) populated clusters 1, 3, 5, 6 and 1 to 5 respectively. While more isoforms were found in Lin-neg cells than tot-BM (28 vs 17 HNRNPM, and 34 vs 8 HNRNPF) the majority in both samples expressed NNIC isoforms (Fig. 6A, B). This is surprising and points to yet unknown novel isoforms in this gene family. PFN1, expressed at higher levels in tot-BM cells is also differentially spliced. Tot-BM cells in clusters 2, 3, 5, and 6 express mainly novel splice variants (NIC) whereas Lin-neg cells express only isoforms matching the reference (FSM) (Fig. 3C, D and 6A).

In summary, our data corroborates that on a single-cell level lineage-negative cells express not only a higher diversity of genes but also more alternatively spliced novel isoforms. From the single-cell analysis, we expand on this finding to reveal that tot-BM cells expressing novel isoforms were mostly located to the CD14+/CD34+ cell cluster 3. In contrast, Lin-neg cells expressing novel isoforms were present in nearly all subpopulations. Further, we discovered that many novel isoforms across all clusters were members of the spliceosome machinery, suggesting a previously unknown functional utility of the observed isoform heterogeneity in human bone marrow progenitor cells.

**Figure 6.**
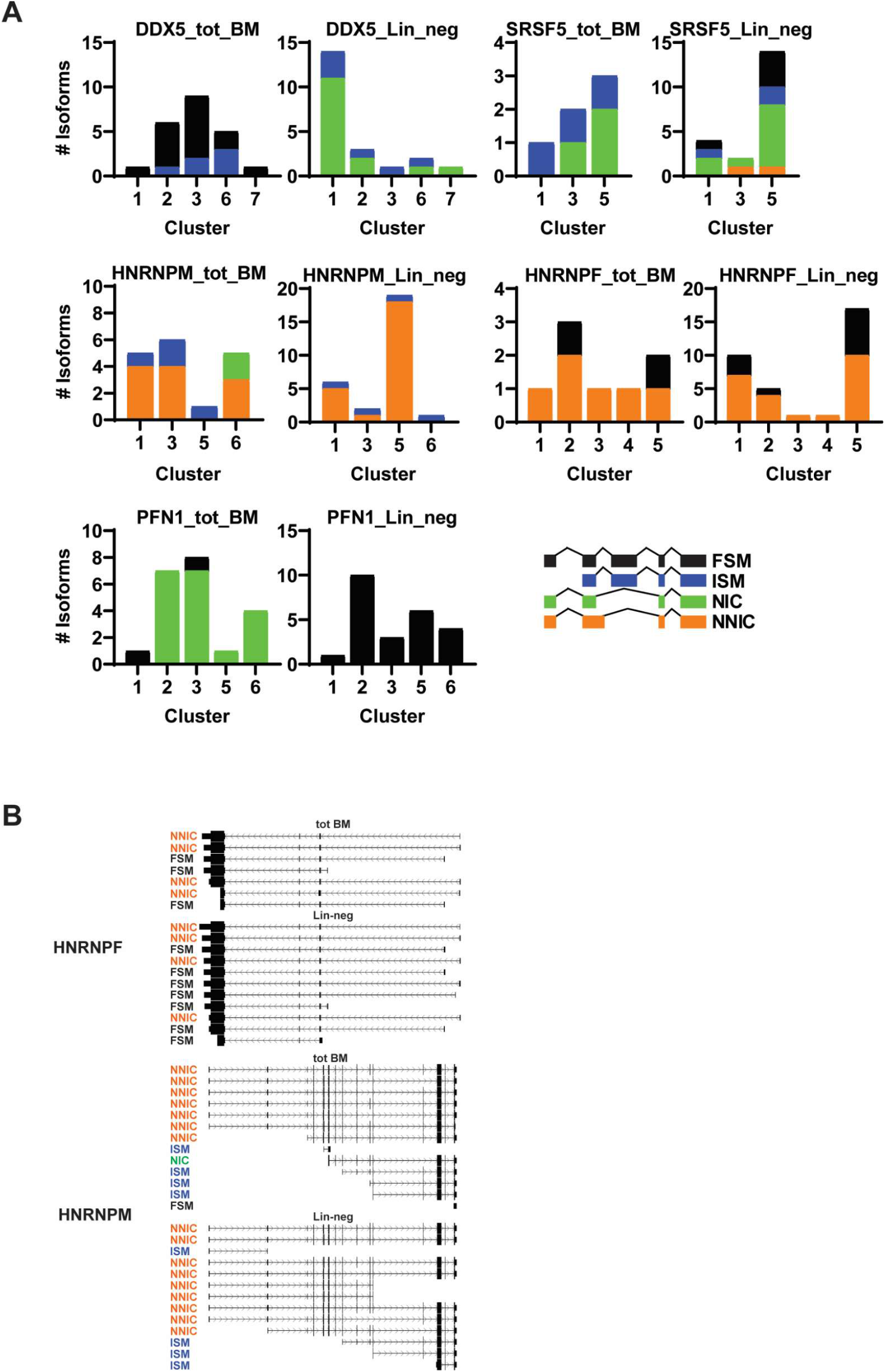
mRNA-processing genes in Lin-neg cells express novel isoforms in most cell type clusters. (**A**) bar graphs showing full-length reads (FL-reads) of isoform categories for DDX5, SRSF5, HNRNPM, HNRNPF, and PFN1. (**B**) USCS genome browser image of full-length isoforms of HNRNPF and HNRNPM. Each isoform is labeled with its respective splicing category. Full Splice Match = FSM, Incomplete Splice Match = ISM, Novel In Catalog = NIC, Novel Not In Catalog = NNIC.

## Discussion

Here we report a high-resolution transcriptional landscape of bone marrow populations both at the level of single-cells and full-length isoforms. We analyzed single-cells from total bone marrow and lineage-negative progenitor subpopulations isolated from the same donor with both short-read and full-length RNA sequencing technology. Previously, we have studied bulk mRNA from the same type of cell populations and found that lineage negative cells express more diverse splice variants including many novel isoforms than the total bone marrow (Deslattes Mays et al. 2019). We now report that this diversity is also reflected at single-cell resolution and not merely a by-product of greater cell type variation being captured in the bulk mRNA homogenate. Lin-neg cells analyzed by short-read sequencing express an expansive variety both in differential expression and in isoform diversity as shown by the number of clusters (9 in Lin-neg vs 5 in tot-BM) and mRNA molecules (133,700 Lin-neg vs 57,300 tot-BM) in a similar number of cells (492 Lin-neg vs 415 tot-BM cells). Full-length transcript analysis indicated an overall increase of novel isoforms. In addition, the Lin-neg cells contained clusters for a wide array of additional progenitor cell clusters such as progenitor cells for granulocytes, neutrophils, basophils, eosinophils, mast cells and B-lymphocytes as well as a subset of CD34-pos stem cells (Table 2 and Fig. 1A). This clustering reflects the greater diversity in Lin-neg cells. After merging both samples in silico and re-clustering, the tot-BM cells populated the additional clusters observed with Lin-neg cells (Fig. 2A). Across the board, Lin-neg cells account for the majority composition in all clusters except for cluster 3 (more mature CD14+ myeloid cells) and cluster 5 (megakaryocytes), where tot-BM cells predominate, reflecting the more mature cell content. The higher diversity of Lin-neg cells indicates the higher plasticity of Lin-neg cells and negates the notion that this population is an amorphous pool of progenitor cells. Rather, our data suggest that the heterogenous Lin-neg progenitor cell subpopulations are poised for different fates.

Also, the single-cell full-length RNA sequencing analysis revealed that individual Lin-neg cells contain more novel isoforms than individual cells in the tot-BM. This extends our previous observation of RNA isoform differences between pools of these cell populations (Deslattes Mays et al. 2019). Interestingly, alternative splicing in Lin-neg cells is found in all subpopulations whereas in tot-BM it is more restricted and predominantly detected in myeloid cells. We were surprised to find spliceosome-associated genes among the genes with the most novel isoforms such as members of the DEAD-box helicases (DDXs), heterogeneous nuclear ribonucleoproteins (HNRNPs), splice regulatory (SR) proteins such as SRSF5, RNase T2 and Splicing Factor 1. In most of the Lin-neg cell clusters these spliceosomal genes including SMG1 were also upregulated relative to tot-BM. It is intriguing that genes involved in mRNA splicing were themselves alternatively spliced as we observed in many cell type clusters of the Lin-neg population. Interestingly, a recent publication (Lee et al. 2018) showed that after knockdowns of HNRNPA1 or DDX5, most alternative splicing events were in genes involved in RNA processing. Among them were SMG7, DDX, SRSF and HNRNP gene family members. Indeed, many recent reviews show that alternative splicing plays a vital role in the self-renewal, pluripotency and lineage specific differentiation of hematopoietic stem cells (Wong et al. 2018; Goldstein et al. 2017; Chen et al. 2014; Li et al. 2021).

Much research has been done on intron retention as a regulatory mechanism for gene expression during hematopoiesis. Intron retention may cause the emergence of a premature termination codon (PTC), which in turn can lead to nonsense-mediated decay (NMD) of mRNA. But NMD is not only an RNA surveillance mechanism in order to degrade mRNAs with nonsense mutations but also serves in an essential regulatory role in post-transcriptional gene expression control in vertebrates (Yi et al. 2020; García-Moreno and Romão 2020; Wong et al. 2018; Lykke-Andersen and Jensen 2015). Finally, previous research has shown that SR proteins are auto-regulated by alternative splicing coupled to nonsense-mediated decay (AS-NMD). SR protein genes comprise conserved regions containing noncoding exons, called ‘’poison exons’’ (PEs), that introduce a PTC and target the mRNA for degradation. We found poison exons expressed predominantly in Lin-neg cells. This finding underscores the role of alternatively spliced poison exons as a potential regulator of gene expression during differentiation of hematopoietic stem cells as described by others (García-Moreno and Romão 2020; Lareau et al. 2007a; Leclair et al. 2020; Jacob and Smith 2017). Overall, our single-cell analysis of bone marrow populations revealed that lineage-negative cells express more diverse splice variants with higher content of novel isoforms. We observed novel isoforms in all Lin-neg cell type clusters and also found them in spliceosome-associated genes. Our findings suggest that isoform diversity by alternative splicing in Lin-neg cells is associated with distinct cell-fate decisions in each cell subpopulation. Future work is needed to explore how alternative splicing is regulating the differentiation of resident bone marrow cells. Especially, investigating the time-dependent expression of novel isoform during maturation of progenitor cells into terminally differentiated cells is of great interest. Furthermore, exploration of the functional activities associated with novel isoforms identified from this analysis could lead to increased understanding of hematopoietic cell differentiation.

## Methods

### Total and lineage negative bone marrow cells

The study was reviewed and considered as “exempt” by the Institutional Review Board of Georgetown University (IRB # 2002-022). All methods were carried out in accordance with relevant guidelines and regulations. Freshly harvested bone marrow tissue was collected from a single discarded healthy human bone marrow collection filter that had been de-identified. mononuclear cells were isolated by Ficoll gradient centrifugation. In order to select for lineage-negative cells, bone marrow mononuclear cells were negatively selected with an antibody cocktail containing antibodies against CD2, CD3, CD5, CD11b, CD11C, CD14, CD16, CD19, CD24, CD61, CD66b, and Glycophorin A (Stemcell Technologies, Vancouver, British Columbia, Canada).

### Single cell cDNA library preparation

Initially 750 of each tot-BM and 750 Lin-neg cells were isolated and then added to the 10x Genomics Chromium controller according to the Chromium Single Cell 3′ Reagent Kits V3 User Guide for a final count of ca. 500 cells. We modified the amplification step (step 2), reducing the cycles from the recommended 14 to 10 in order to capture more rare transcripts. The amplified cDNA library was then spit. 25% was used for Illumina short-read library preparation and 75% for the PacBio library preparation.

### Short-read RNA-seq

For the short-read library preparation the chromium user guide was continued and the final fragmented cDNA libraries were sent to Novagene (Sacramento, CA) and sequenced on an Illumina Hiseq 4000 with a depth of 100 million paired reads per sample and a read length of 150nt. The resulting raw reads (FASTQ) were then de-multiplexed using cellranger software (10xgenomics). The reads were filtered for highly diverse UMI in order to remove bulk non-single cell-derived transcripts. The reads were then clustered with the Cell-Loupe browser (10xGenomics) to ascertain the quality of samples. The resulting non-normalized data were then fed into the Seurat pipeline (Stuart et al. 2019) for more detailed analysis.

### Full-length RNA-seq

The cDNA libraries containing sample index, UMI, and the cell barcode from step 2 of the 10xpipeline was sent to University of Maryland’s Institute for Genome Sciences for full-length sequencing on the PacBio Sequel 2 platform. The two sample libraries were pooled with two other libraries and run for 30 hours. The resulting circular consensus (CSS) reads were demultiplexed, filtered for consensus within 3 passes and PacBio’s Isoseq3 (https://github.com/PacificBiosciences/IsoSeq) pipeline was followed with modifications for single cell data (Cupcake by Elizabeth Tseng https://github.com/Magdoll/cDNA_Cupcake, SQANTI3: https://github.com/ConesaLab/SQANTI3). In short 3’ and 5’ primers were detected and removed (Lima), then UMIs and Cell barcodes were detected (clip_out_UMI_cellBC.py), finally the polyA tail was detected and artificial concatemers were removed (isoseq3 refine). Then the reads were aligned to the genome (minmap2) and unique transcripts were collapsed (collapse_isoforms_by_sam.py). The reads were then annotated with sqanti3_qc.py, artefacts were filtered out (sqanti3_RulesFilter.py), and the high-quality transcripts were combined with the UMI and cell barcodes for the final report (collate_FLNC_gene_info.py, UMI_BC_error_correct.py)

### Single cell cluster analysis (Seurat, (Stuart et al. 2019))

Single cell analysis followed vignettes posted in github: https://github.com/satijalab/seurat. Briefly, count matrix output from Cellranger (10xgenomics) was read into Seurat software on R. Quality control of the count matrix was performed by filtering out single cells with more than 25% mitochondrial mRNA content and less than 200 UMIs as those represent empty droplets. The UMI counts were normalized, log transformed and scaled by linear transformation to equalize mean and variance across genes. Principal component analysis and determination of dimensionality of the data was performed and the data was visualized by clustering the cells using non-linear dimensional reduction (tSNE or UMAP). In order to annotate the cell types of each cluster differentially expressed features were used as find biomarker. For differential analysis the Seurat objects for each sample (tot-BM and Lin-neg) were merged and the differential gene expression analysis was repeated. Barcodes from each cluster were identified and used to select and identify full-length reads for isoform analysis.

## Code availability

The code will be deposited as a Nextflow pipeline. Custom code is available upon request.

## Data access

The raw and processed data is available from the Gene Expression Omnibus repository (GSE181160).

## Competing interest

None of the authors have any competing interests

## Acknowledgments

We are grateful to the leadership and staff of the bone marrow donor program at the Georgetown University Hospital for their continued support. We acknowledge support by the Genomics and Epigenomics Shared Resources of the Lombardi Comprehensive Cancer Center.

## Notes

### Competing Interest Statement

The authors have declared no competing interest.

## References

Arzalluz-Luque Á, Conesa A. 2018. Single-cell RNAseq for the study of isoforms—how is that possible? Genome Biol 19: 110.

Boise LH, González-García M, Postema CE, Ding L, Lindsten T, Turka LA, Mao X, Nuñez G, Thompson CB. 1993. bcl-x, a bcl-2-related gene that functions as a dominant regulator of apoptotic cell death. Cell 74: 597–608.

Bourgeois CF, Mortreux F, Auboeuf D. 2016. The multiple functions of RNA helicases as drivers and regulators of gene expression. Nat Rev Mol Cell Bio 17: 426–438.

Byrne A, Beaudin AE, Olsen HE, Jain M, Cole C, Palmer T, DuBois RM, Forsberg EC, Akeson M, Vollmers C. 2017. Nanopore long-read RNAseq reveals widespread transcriptional variation among the surface receptors of individual B cells. Nat Commun 8: 16027.

Cheng J, Zhou T, Liu C, Shapiro J, Brauer M, Kiefer M, Barr P, Mountz J. 1994. Protection from Fas-mediated apoptosis by a soluble form of the Fas molecule. Science 263: 1759–1762.

Chen J, Crutchley J, Zhang D, Owzar K, Kastan MB. 2017. Identification of a DNA Damage– Induced Alternative Splicing Pathway That Regulates p53 and Cellular Senescence Markers. Cancer Discov 7: 766–781.

Chen L, Kostadima M, Martens JHA, Canu G, Garcia SP, Turro E, Downes K, Macaulay IC, Bielczyk-Maczynska E, Coe S, et al. 2014. Transcriptional diversity during lineage commitment of human blood progenitors. Science 345: 1251033.

Chen M, Zhang J, Manley JL. 2010. Turning on a Fuel Switch of Cancer: hnRNP Proteins Regulate Alternative Splicing of Pyruvate Kinase mRNA. Cancer Res 70: 8977–8980.

Consortium IHGS. 2004. Finishing the euchromatic sequence of the human genome. Nature 431: 931–945.

García-Moreno JF, Romão L. 2020. Perspective in Alternative Splicing Coupled to Nonsense-Mediated mRNA Decay. Int J Mol Sci 21: 9424.

Goldstein O, Meyer K, Greenshpan Y, Bujanover N, Feigin M, Ner-Gaon H, Shay T, Gazit R. 2017. Mapping Whole-Transcriptome Splicing in Mouse Hematopoietic Stem Cells. Stem Cell Rep 8: 163–176.

Gupta I, Collier PG, Haase B, Mahfouz A, Joglekar A, Floyd T, Koopmans F, Barres B, Smit AB, Sloan SA, et al. 2018. Single-cell isoform RNA sequencing characterizes isoforms in thousands of cerebellar cells. Nat Biotechnol 36: 1197–1202.

Hardwick SA, Joglekar A, Flicek P, Frankish A, Tilgner HU. 2019. Getting the Entire Message: Progress in Isoform Sequencing. Frontiers Genetics 10: 709.

Jacob AG, Smith CWJ. 2017. Intron retention as a component of regulated gene expression programs. Hum Genet 136: 1043–1057.

Karlsson K, Lönnerberg P, Linnarsson S. 2017. Alternative TSSs are co-regulated in single cells in the mouse brain. Mol Syst Biol 13: 930.

Lareau LF, Brooks AN, Soergel DAW, Meng Q, Brenner SE. 2007a. Alternative Splicing in the Postgenomic Era. Adv Exp Med Biol 623: 190–211.

Lareau LF, Inada M, Green RE, Wengrod JC, Brenner SE. 2007b. Unproductive splicing of SR genes associated with highly conserved and ultraconserved DNA elements. Nature 446: 926–929.

Leclair NK, Brugiolo M, Urbanski L, Lawson SC, Thakar K, Yurieva M, George J, Hinson JT, Cheng A, Graveley BR, et al. 2020. Poison Exon Splicing Regulates a Coordinated Network of SR Protein Expression during Differentiation and Tumorigenesis. Mol Cell 80: 648–665.e9.

Lee YJ, Wang Q, Rio DC. 2018. Coordinate regulation of alternative pre-mRNA splicing events by the human RNA chaperone proteins hnRNPA1 and DDX5. Gene Dev 32: 1060–1074.

Li Y, Wang D, Wang H, Huang X, Wen Y, Wang B, Xu C, Gao J, Liu J, Tong J, et al. 2021. A splicing factor switch controls hematopoietic lineage specification of pluripotent stem cells. Embo Rep 22: e50535.

Lykke-Andersen S, Jensen TH. 2015. Nonsense-mediated mRNA decay: an intricate machinery that shapes transcriptomes. Nat Rev Mol Cell Bio 16: 665–677.

Deslattes Mays A, Schmidt M, Graham G, Tseng E, Baybayan P, Sebra R, Sanda M, Mazarati J-B, Riegel A, Wellstein A. 2019. Single-Molecule Real-Time (SMRT) Full-Length RNA-Sequencing Reveals Novel and Distinct mRNA Isoforms in Human Bone Marrow Cell Subpopulations. Genes-basel 10: 253.

Nowak DG, Woolard J, Amin EM, Konopatskaya O, Saleem MA, Churchill AJ, Ladomery MR, Harper SJ, Bates DO. 2008. Expression of pro- and anti-angiogenic isoforms of VEGF is differentially regulated by splicing and growth factors. J Cell Sci 121: 3487–3495.

Pellin D, Loperfido M, Baricordi C, Wolock SL, Montepeloso A, Weinberg OK, Biffi A, Klein AM, Biasco L. 2019. A comprehensive single cell transcriptional landscape of human hematopoietic progenitors. Nat Commun 10: 2395.

Powers KT, Szeto J-YA, Schaffitzel C. 2020. New insights into no-go, non-stop and nonsense-mediated mRNA decay complexes. Curr Opin Struc Biol 65: 110–118.

Stuart T, Butler A, Hoffman P, Hafemeister C, Papalexi E, Mauck WM, Hao Y, Stoeckius M, Smibert P, Satija R. 2019. Comprehensive Integration of Single-Cell Data. Cell 177: 1888–1902.e21.

Tardaguila M, Fuente L de la, Marti C, Pereira C, Pardo-Palacios FJ, Risco H del, Ferrell M, Mellado M, Macchietto M, Verheggen K, et al. 2018. SQANTI: extensive characterization of long-read transcript sequences for quality control in full-length transcriptome identification and quantification. Genome Res 28: 396–411.

Twyffels L, Gueydan C, Kruys V. 2011. Shuttling SR proteins: more than splicing factors. Febs J 278: 3246–3255.

Velten L, Haas SF, Raffel S, Blaszkiewicz S, Islam S, Hennig BP, Hirche C, Lutz C, Buss EC, Nowak D, et al. 2017. Human haematopoietic stem cell lineage commitment is a continuous process. Nat Cell Biol 19: 271–281.

Vieler M, Sanyal S. 2018. p53 Isoforms and Their Implications in Cancer. Cancers 10: 288.

Wong ACH, Rasko JEJ, Wong JJ-L. 2018. We skip to work: alternative splicing in normal and malignant myelopoiesis. Leukemia 32: 1081–1093.

Yi Z, Sanjeev M, Singh G. 2020. The Branched Nature of the Nonsense-Mediated mRNA Decay Pathway. Trends Genet 37: 143–159.

