## supplemental material for "Full-length, single-cell RNA-sequencing of human bone marrow subpopulations reveals hidden complexity"

**Supplemental data:**

Suppl. Figure S1

Combined tot-BM and Lin-neg marker genes for each cluster.

Suppl. Figure S2

Spliceosome-associated transcripts contain poison exons

Suppl. Table S1

Tot-BM marker genes for each cluster

Suppl. Table S2

Lin-neg marker genes for each cluster.

Suppl. Table S3

Combined tot-BM and Lin-neg marker genes for each cluster.

### Figure S1

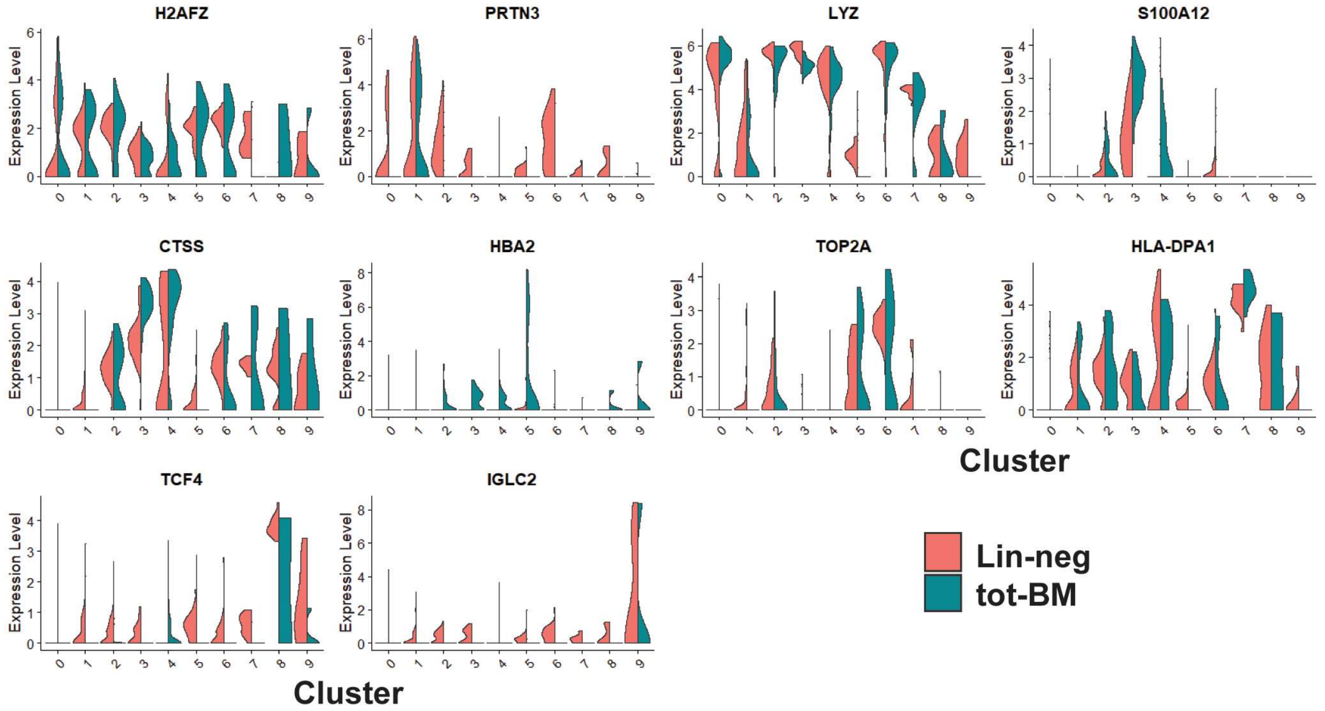


**Suppl. Figure S1.** Top marker gene for each cell type cluster. Gene names with their expression levels and p-values are in Suppl. Table S3

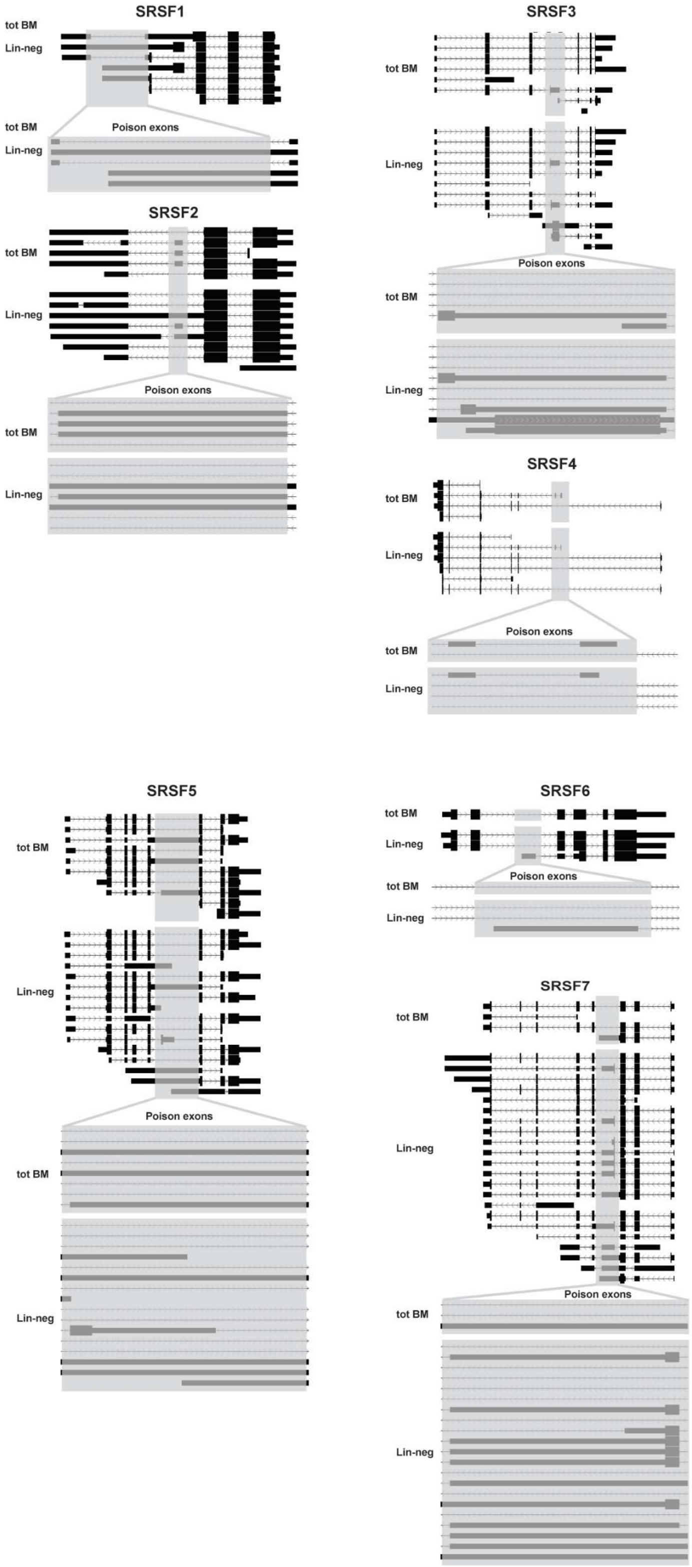


**Suppl. Figure S2.** Spliceosome-associated transcripts contain poison exons. Isoforms detected by full-length sequencing for SRSF1 to 7. Poison exons are highlighted in grey. Analysis from the UCSC browser

**Suppl. Table S1 Tot-BM marker genes for each cluster.** Statistics calculated comparing expression in cluster-specific cells vs cells in all other clusters. Columns are: Log-fold change, multiple test p-value, Bonferroni adjusted p-value, Percentage of positive cells in spec. cluster, Percentage of positive cells in all other clusters,

| gene | name | cluster | Fold change (log) | p-value | p-value (adj) | pct. spec custer | pct. other clusters |
|---|---|---|---|---|---|---|---|
| **IGKC** | major histocompatibility complex, class II, DM alpha | 0 | 1.7065626 | 1.71E-06 | 2.01E-02 | 0.038 | 0.240 |
| **MPO** | myeloperoxidase | 0 | 0.85762519 | 0.00041 | 1.00E+00 | 0.583 | 0.548 |
| **HEXB** | hypophosphatemic bone disease | 0 | 0.832369 | 0.00039 | 1.00E+00 | 0.129 | 0.343 |
| **HLA-DMA** | hexosaminidase subunit beta | 0 | 0.75546393 | 0.00554 | 1.00E+00 | 0.152 | 0.329 |
| **S100A12** | S100 calcium binding protein A12 | 1 | 2.45585861 | 2.25E-64 | 2.64E-60 | 0.886 | 0.058 |
| **CTSS** | cathepsin S | 1 | 2.37620171 | 2.63E-61 | 3.08E-57 | 1.000 | 0.174 |
| **FCN1** | ficolin 1 | 1 | 2.06343714 | 7.21E-61 | 8.44E-57 | 0.981 | 0.132 |
| **RGS2** | regulator of G-protein signaling 2 | 1 | 2.03424637 | 1.86E-66 | 2.18E-62 | 0.943 | 0.068 |
| **HBB** | hemoglobin subunit beta | 2 | 2.17273693 | 8.34E-06 | 9.76E-02 | 0.485 | 0.371 |
| **HBD** | hemoglobin subunit delta | 2 | 2.00152579 | 5.75E-12 | 6.73E-08 | 0.237 | 0.028 |
| **AHSP** | alpha hemoglobin stabilizing protein | 2 | 1.99110243 | 1.08E-06 | 1.26E-02 | 0.247 | 0.082 |
| **PRDX2** | peroxiredoxin 2 | 2 | 1.7008398 | 1.56E-10 | 1.83E-06 | 0.443 | 0.186 |
| **TOP2A** | topoisomerase (DNA) II alpha | 3 | 1.43339091 | 1.12E-06 | 1.31E-02 | 0.359 | 0.140 |
| **CENPF** | centromere protein F | 3 | 1.35619246 | 4.09E-07 | 4.79E-03 | 0.359 | 0.131 |
| **MKI67** | lysozyme | 3 | 1.31678345 | 5.33E-07 | 6.24E-03 | 0.375 | 0.145 |
| **UBE2C** | ubiquitin conjugating enzyme E2 C | 3 | 1.30342572 | 1.43E-05 | 1.67E-01 | 0.219 | 0.063 |
| **NRGN** | neurogranin | 4 | 0.81717354 | 3.61E-14 | 4.23E-10 | 1.000 | 0.226 |
| **MKI67** | marker of proliferation Ki-67 | 4 | 0.81578158 | 1.64E-12 | 1.92E-08 | 0.882 | 0.151 |
| **EREG** | epiregulin | 4 | 0.79205986 | 3.33E-06 | 3.90E-02 | 0.706 | 0.216 |
| **LYZ** | immunoglobulin kappa constant | 4 | 0.78636841 | 5.29E-06 | 6.20E-02 | 1.000 | 0.731 |

**Suppl. Table S2 Lin-neg marker genes for each cluster.** Statistics calculated comparing expression in cluster-specific cells vs cells in all other clusters. Columns are: Log-fold change, multiple test p-value, Bonferroni adjusted p-value, Percentage of positive cells in spec. cluster, Percentage of positive cells in all other clusters,

| gene | name | cluster | Fold change (log) | p-value | p-value (adj) | pct. spec custer | pct. other clusters |
|---|---|---|---|---|---|---|---|
| **CTSG** | cathepsin G | 0 | 1.242 | 4.80E-03 | 1.00E+00 | 0.285 | 0.575 |
| **DEFA4** | defensin alpha 4 | 0 | 1.201 | 3.21E-03 | 1.00E+00 | 0.032 | 0.117 |
| **ZNF90** | zinc finger protein 90 | 0 | 1.182 | 5.71E-04 | 1.00E+00 | 0.032 | 0.138 |
| **HIST3H2A** | histone cluster 3 H2A | 0 | 0.941 | 3.21E-03 | 1.00E+00 | 0.032 | 0.117 |
| **S100A8** | S100 calcium binding protein A8 | 1 | 0.998 | 3.83E-22 | 5.43E-18 | 0.933 | 0.481 |
| **LYZ** | lysozyme | 1 | 0.963 | 6.20E-27 | 8.79E-23 | 0.983 | 0.817 |
| **CXCL8** | C-X-C motif chemokine ligand 8 | 1 | 0.952 | 5.89E-24 | 8.36E-20 | 0.592 | 0.145 |
| **SLC2A3** | solute carrier family 2 member 3 | 1 | 0.797 | 4.36E-35 | 6.19E-31 | 0.800 | 0.204 |
| **FCN1** | ficolin 1 | 2 | 1.953 | 6.61E-08 | 9.38E-04 | 0.554 | 0.328 |
| **CTSS** | cathepsin S | 2 | 1.719 | 1.58E-03 | 1.00E+00 | 0.536 | 0.571 |
| **S100A9** | S100 calcium binding protein A9 | 2 | 1.352 | 1.24E-09 | 1.76E-05 | 0.786 | 0.622 |
| **VCAN** | versican | 2 | 1.302 | 1.09E-04 | 1.00E+00 | 0.500 | 0.346 |
| **IGLL1** | immunoglobulin lambda like polypeptide 1 | 3 | 1.415 | 4.29E-27 | 6.08E-23 | 0.830 | 0.175 |
| **SPINK2** | serine peptidase inhibitor, Kazal type 2 | 3 | 1.103 | 7.58E-34 | 1.08E-29 | 0.745 | 0.088 |
| **C1QTNF4** | C1q and tumor necrosis factor related protein 4 | 3 | 1.077 | 1.33E-43 | 1.88E-39 | 0.894 | 0.097 |
| **PRSS57** | protease, serine 57 | 3 | 0.961 | 1.29E-27 | 1.83E-23 | 0.957 | 0.245 |
| **TOP2A** | topoisomerase | 4 | 1.554 | 5.27E-29 | 7.48E-25 | 1.000 | 0.300 |
| **CLC** | Charcot-Leyden crystal galectin | 4 | 1.405 | 5.65E-05 | 8.01E-01 | 0.190 | 0.042 |
| **MKI67** | marker of proliferation Ki-67 | 4 | 1.399 | 6.37E-27 | 9.04E-23 | 0.976 | 0.324 |
| **CENPF** | centromere protein F | 4 | 1.366 | 3.65E-24 | 5.18E-20 | 0.952 | 0.298 |
| **HBB** | hemoglobin subunit beta | 5 | 3.089 | 2.26E-20 | 3.21E-16 | 1.000 | 0.348 |
| **AHSP** | alpha hemoglobin stabilizing protein | 5 | 1.940 | 1.83E-39 | 2.59E-35 | 0.897 | 0.084 |
| **PRDX2** | peroxiredoxin 2 | 5 | 1.886 | 9.85E-23 | 1.40E-18 | 1.000 | 0.326 |
| **HBA1** | hemoglobin subunit alpha 1 | 5 | 1.840 | 9.97E-35 | 1.41E-30 | 0.690 | 0.045 |
| **HLA-DPB1** | major histocompatibility complex, class II, DP beta 1 | 6 | 2.567 | 2.58E-11 | 3.66E-07 | 1.000 | 0.545 |
| **HLA-DQA1** | major histocompatibility complex, class II, DQ alpha 1 | 6 | 2.432 | 3.99E-14 | 5.67E-10 | 1.000 | 0.304 |
| **HLA-DPA1** | major histocompatibility complex, class II, DP alpha 1 | 6 | 2.412 | 3.98E-11 | 5.65E-07 | 1.000 | 0.570 |
| **ID2** | inhibitor of DNA binding 2, HLH protein | 6 | 2.317 | 3.72E-16 | 5.27E-12 | 1.000 | 0.260 |
| **TCF4** | transcription factor 4 | 7 | 3.115 | 6.23E-13 | 8.84E-09 | 1.000 | 0.374 |
| **GZMB** | granzyme B | 7 | 2.712 | 2.78E-32 | 3.94E-28 | 0.929 | 0.065 |
| **HIST1H2BG** | histone cluster 1 H2B family member g | 7 | 2.620 | 1.70E-26 | 2.42E-22 | 0.929 | 0.088 |
| **CCDC50** | coiled-coil domain containing 50 | 7 | 2.594 | 4.03E-15 | 5.72E-11 | 1.000 | 0.266 |
| **IGHA1** | immunoglobulin heavy constant alpha 1 | 8 | 4.840 | 4.83E-04 | 1.00E+00 | 0.818 | 0.516 |
| **IGKC** | immunoglobulin kappa constant | 8 | 4.496 | 6.32E-05 | 8.96E-01 | 1.000 | 0.655 |
| **IGLC3** | immunoglobulin lambda constant 3 | 8 | 4.474 | 1.56E-04 | 1.00E+00 | 0.364 | 0.075 |
| **IGHG1** | immunoglobulin heavy constant gamma 1 | 8 | 3.818 | 3.08E-09 | 4.37E-05 | 1.000 | 0.356 |

**Table S3, Combined tot-BM and Lin-neg marker genes for each cluster. .** Statistics calculated comparing expression in cluster-specific cells vs cells in all other clusters. Columns are: Log-fold change, multiple test p-value, Bonferroni adjusted p-value, Percentage of positive cells in spec. cluster, Percentage of positive cells in all other clusters,

| gene | name | cluster | Fold change (log) | p-value | p-value (adj) | pct. spec custer | pct. other clusters |
|---|---|---|---|---|---|---|---|
| **H2AFZ** | H2A histone family member Z | 0 | 0.880 | 8.16E-04 | 1.00E+00 | 0.352 | 0.703 |
| **EIF5** | eukaryotic translation initiation factor 5 | 0 | 0.822 | 8.43E-16 | 1.21E-11 | 0.146 | 0.559 |
| **CFD** | complement factor D | 0 | 0.698 | 7.09E-06 | 1.02E-01 | 0.188 | 0.441 |
| **UBB** | ubiquitin B | 0 | 0.608 | 3.21E-07 | 4.59E-03 | 0.272 | 0.657 |
| **PRTN3** | proteinase 3 | 1 | 1.978 | 3.47E-11 | 4.96E-07 | 0.517 | 0.333 |
| **CTSG** | cathepsin G | 1 | 1.885 | 1.42E-06 | 2.04E-02 | 0.453 | 0.309 |
| **ELANE** | elastase, neutrophil expressed | 1 | 1.794 | 3.73E-07 | 5.33E-03 | 0.535 | 0.405 |
| **IGLL1** | immunoglobulin lambda like polypeptide 1 | 1 | 1.632 | 1.19E-37 | 1.70E-33 | 0.465 | 0.093 |
| **LYZ** | lysozyme | 2 | 0.735 | 8.76E-24 | 1.25E-19 | 0.992 | 0.776 |
| **PCNA** | proliferating cell nuclear antigen | 2 | 0.702 | 2.51E-38 | 3.60E-34 | 0.867 | 0.271 |
| **CDCA7** | cell division cycle associated 7 | 2 | 0.612 | 1.53E-52 | 2.19E-48 | 0.725 | 0.130 |
| **HELLS** | helicase, lymphoid-specific | 2 | 0.568 | 1.96E-43 | 2.81E-39 | 0.775 | 0.183 |
| **S100A12** | S100 calcium binding protein A12 | 3 | 2.216 | 4.21E-82 | 6.02E-78 | 0.928 | 0.143 |
| **SLC2A3** | solute carrier family 2 member 3 | 3 | 1.847 | 3.47E-67 | 4.97E-63 | 0.990 | 0.259 |
| **S100A8** | S100 calcium binding protein A8 | 3 | 1.816 | 8.41E-51 | 1.20E-46 | 1.000 | 0.498 |
| **S100A9** | S100 calcium binding protein A9 | 3 | 1.813 | 1.05E-50 | 1.50E-46 | 1.000 | 0.556 |
| **CTSS** | cathepsin S | 4 | 1.525 | 1.59E-14 | 2.28E-10 | 0.677 | 0.461 |
| **FCN1** | ficolin 1 | 4 | 1.387 | 1.09E-21 | 1.56E-17 | 0.731 | 0.307 |
| **POU2F2** | POU class 2 homeobox 2 | 4 | 1.314 | 2.48E-08 | 3.54E-04 | 0.473 | 0.290 |
| **HLA-DQB1** | major histocompatibility complex, class II, DQ beta 1 | 4 | 1.298 | 9.35E-14 | 1.34E-09 | 0.710 | 0.471 |
| **HBA2** | hemoglobin subunit alpha 2 | 5 | 4.336 | 2.40E-06 | 3.43E-02 | 0.300 | 0.127 |
| **HBB** | hemoglobin subunit beta | 5 | 4.129 | 1.52E-38 | 2.17E-34 | 0.875 | 0.345 |
| **HBA1** | hemoglobin subunit alpha 1 | 5 | 4.033 | 2.44E-21 | 3.49E-17 | 0.525 | 0.139 |
| **AHSP** | alpha hemoglobin stabilizing protein | 5 | 2.410 | 4.38E-55 | 6.26E-51 | 0.650 | 0.076 |
| **TOP2A** | topoisomerase | 6 | 1.582 | 3.49E-32 | 4.99E-28 | 0.792 | 0.230 |
| **MKI67** | marker of proliferation Ki-67 | 6 | 1.447 | 1.93E-34 | 2.77E-30 | 0.819 | 0.243 |
| **CENPE** | centromere protein E | 6 | 1.399 | 3.07E-39 | 4.40E-35 | 0.708 | 0.132 |
| **CENPF** | centromere protein F | 6 | 1.342 | 1.33E-28 | 1.90E-24 | 0.764 | 0.225 |
| **HLA-DPA1** | major histocompatibility complex, class II, DP alpha 1 | 7 | 2.663 | 2.05E-20 | 2.93E-16 | 1.000 | 0.490 |
| **HLA-DQA1** | major histocompatibility complex, class II, DQ alpha 1 | 7 | 2.597 | 1.98E-22 | 2.83E-18 | 0.889 | 0.227 |
| **HLA-DPB1** | major histocompatibility complex, class II, DP beta 1 | 7 | 2.524 | 1.21E-16 | 1.73E-12 | 0.926 | 0.452 |
| **HLA-DRB1** | major histocompatibility complex, class II, DR beta 1 | 7 | 2.246 | 7.19E-19 | 1.03E-14 | 1.000 | 0.555 |
| **TCF4** | transcription factor 4 | 8 | 3.093 | 4.06E-15 | 5.81E-11 | 0.889 | 0.268 |
| **CCDC50** | coiled-coil domain containing 50 | 8 | 2.615 | 1.67E-18 | 2.39E-14 | 0.889 | 0.196 |
| **IRF4** | interferon regulatory factor 4 | 8 | 2.474 | 1.07E-49 | 1.53E-45 | 0.778 | 0.035 |
| **HIST1H2BG** | histone cluster 1 H2B family member g | 8 | 2.448 | 4.12E-30 | 5.89E-26 | 0.722 | 0.058 |
| **IGLC2** | immunoglobulin lambda constant 2 | 9 | 6.099 | 2.43E-03 | 1.00E+00 | 0.467 | 0.215 |
| **IGKC** | immunoglobulin kappa constant | 9 | 5.356 | 2.46E-08 | 3.52E-04 | 0.933 | 0.432 |
| **IGHA1** | immunoglobulin heavy constant alpha 1 | 9 | 5.013 | 1.63E-06 | 2.33E-02 | 0.733 | 0.280 |
| **IGHG1** | immunoglobulin heavy constant gamma 1 | 9 | 4.516 | 1.37E-15 | 1.96E-11 | 0.933 | 0.211 |
